# Population differences in the length and early-life dynamics of telomeres among European pied flycatchers

**DOI:** 10.1101/2021.03.16.435615

**Authors:** Tiia Kärkkäinen, Toni Laaksonen, Malcolm Burgess, Alejandro Cantarero, Jesús Martínez-Padilla, Jaime Potti, Juan Moreno, Robert L. Thomson, Vallo Tilgar, Antoine Stier

## Abstract

Telomere length and shortening rate are increasingly used as biomarkers for long-term costs in ecological and evolutionary studies because of their relationships with survival and fitness. Both early-life conditions and growth, and later-life stressors can create variation in telomere shortening rate. Studies on between-population telomere length and dynamics are scarce, despite the expectation that populations exposed to varying environmental constraints would present divergent telomere length patterns. The pied flycatcher (*Ficedula hypoleuca*) is a passerine bird breeding across Eurasia (from Spain to western Siberia) and migrating through the Iberian Peninsula to spend the non-breeding period in sub-Saharan Africa. Thus, different populations show marked differences in migration distance. We studied the large-scale variation of telomere length and early-life dynamics in the pied flycatcher by comparing six European populations across a north-south gradient (Finland, Estonia, England, and Spain) predicting negative effect of migration distance on adult telomere length, and of nestling growth on nestling telomere dynamics. There were clear population differences in telomere length, with English birds from mid-latitudes having the longest telomeres. Telomere length did not thus show consistent latitudinal variation and was not linearly linked to differences in migration distance. Early-life telomere shortening rate tended to vary between populations. Fast growth was associated with shorter telomeres in the early life, but faster nestling growth affected telomeres more negatively in northern than southern populations. While the sources of between-population differences in telomere-related biology remain to be more intensively studied, our study illustrates the need to expand telomere studies at the between-population level.

## Introduction

Telomeres, the capping structures of linear chromosomes, have a crucial role in maintaining genomic integrity and cell viability (Blackburn, 1991; Blackburn, Epel, & Lin, 2015). They shorten with cell divisions and the shortening can be accentuated by cellular and external stressors, such as oxidative stress or substantially high energy demands (Casagrande & Hau, 2019; Levy, Allsopp, Futcher, Greider, & Harley, 1992; Reichert & Stier, 2017). As short telomeres are associated with ageing phenotypes (Campisi, Kim, Lim, & Rubio, 2001), telomere length is increasingly used as a biomarker of ageing to predict survival and fitness (Monaghan, Eisenberg, Harrington, & Nussey, 2018). To date in wild populations, telomere length has been associated with past stress exposure (Chatelain, Drobniak, & Szulkin, 2020), individual quality (Angelier, Weimerskirch, Barbraud, & Chastel, 2019), fitness (Eastwood et al., 2019) and overall mortality (Wilbourn et al., 2018), suggesting usefulness of telomere length as a biomarker for long-term costs in wild animals.

Most telomere shortening happens during early life growth (Spurgin et al., 2018; Stier, Metcalfe, & Monaghan, 2020), and fast growth has been suggested to accelerate telomere shortening (Monaghan & Ozanne, 2018). Early-life conditions, with the associated hormone levels (Casagrande et al., 2020; Stier, Hsu, et al., 2020), competition (Cram, Monaghan, Gillespie, & Clutton-Brock, 2017; Young et al., 2017), and nutrition deficiency (Nettle et al., 2017) can affect individual telomere length trajectories and thus could promote individual differences in longevity. Later-life stressors, such as predation risk (Kärkkäinen et al., 2019), parasitic infections (Asghar et al., 2015), low prey abundance (Spurgin et al., 2018), reproductive effort (Bauch, Gatt, Granadeiro, Verhulst, & Catry, 2020; López-Arrabé et al., 2018; Sudyka, Arct, Drobniak, Gustafsson, & Cichoń, 2019) and migration (Bauer, Heidinger, Ketterson, & Greives, 2016) can create further between-individual differences in telomere length. Telomere length and dynamics have also been associated with genetic polymorphism (Eisenberg, 2019; Karell, Bensch, Ahola, & Asghar, 2017).

While within-population telomere length patterns have been widely examined (*i.e.* most of the examples cited above), studies on among-population telomere length and dynamics are still scarce (Burraco, Lucas, & Salmón, 2021). Species distributed over vast latitudinal gradients face different and variable environmental conditions, for example in respect to temperature and seasonality (Willig, Kaufman, & Stevens, 2003). Indeed, life-histories across species vary often in a latitudinal manner with high latitude species more likely exhibiting a faster pace of life characterized by higher basal metabolic rate and lower adult survival than low latitude ones (Muñoz, Kéry, Martins, & Ferraz, 2018; Wikelski, Spinney, Schelsky, Scheuerlein, & Gwinner, 2003). Consequently, many species-specific life-history traits and strategies, *e.g.*, clutch size, parental investment, and juvenile growth rate vary in a latitudinal gradient (McNamara, Barta, Wikelski, & Houston, 2008). Thus through possible differences in the pace of life, latitudinal variation might ultimately influence also telomere dynamics (Angelier, Costantini, Blévin, & Chastel, 2018; Giraudeau, Angelier, & Sepp, 2019). While across species, fast paced and shorter lived species have longer telomeres (Pepke, Ringsby, & Eisenberg, 2021) and faster telomere attrition (Dantzer & Fletcher, 2015), the opposite pattern is expected at the within-species level, with populations of high latitudes being predicted to have shorter telomeres and faster attrition than populations of low latitudes (Giraudeau et al., 2019). Accordingly, telomere length has been shown to decrease at higher latitudes in American black bears (*Ursus americanus*) (Kirby, Alldredge, & Pauli, 2017). Similar to latitudinal gradients, increasing elevation can change environmental factors (Hille & Cooper, 2015; Willig et al., 2003). For example, nestlings from two tit species (*Parus* spp.) showed faster telomere shortening in populations breeding at higher altitudes than in populations breeding at lower altitudes (Stier et al., 2016). Telomere length has been associated with geography and ethnicity also in humans (Hunt et al., 2020; Ly et al., 2019). Therefore, populations or subpopulations of species facing specific energetic demands and environmental stressors may display divergent patterns of telomere length and dynamics, and ultimately ageing rates (Ibáñez-Álamo et al., 2018). Knowledge of the mechanisms driving individuals’ telomere length trajectories across populations could help in understanding life history evolution in different environments and the resilience of populations to environmental change, as short telomeres could also be indicative of local extinction risk (Dupoué et al., 2017). In addition, potential differences in telomere length between populations within a species question the use of ‘species’ data in meta-analyses and comparative studies when there are data from only one population.

Our study species, the pied flycatcher (*Ficedula hypoleuca*), is a small, insectivorous migratory passerine that breeds over a large area of Eurasia from Spain to western Siberia in a wide range of woodland habitats, from high altitude forests to temperate deciduous and boreal coniferous forests (Lundberg & Alatalo, 1992). In the autumn, pied flycatchers from across the breeding range migrate through the Iberian peninsula to spend the non-breeding period in sub-Saharan Africa (Chernetsov, Kishkinev, Gashkov, Kosarev, & Bolshakov, 2008; Lundberg & Alatalo, 1992; Ouwehand et al., 2016). Consequently, different pied flycatcher populations experience marked differences in the distance (and hence duration) of their migration. Migratory flight can increase metabolic rate (Kvist & Lindström, 2001), which in turn might accelerate telomere shortening either through increased oxidative stress (Reichert & Stier, 2017) or through metabolic adjustments (Casagrande & Hau, 2019). Furthermore, migrant species have faster pace of life than resident species (Soriano-Redondo, Gutiérrez, Hodgson, & Bearhop, 2020), and it is possible that within species longer migrations could result in faster pace of life due to increases in used energy and risks related to migration, such as elevated mortality (Sillett & Holmes, 2002). Thus, pied flycatchers with a longer migration distance (northern populations) might exhibit shorter telomeres than those with a shorter migration (southern populations). Accordingly, there is evidence that pied flycatcher females breeding in Spain, in the southern part of the breeding range and with the shortest migration, show higher adult survival, natal recruitment rate and delayed onset of reproductive ageing compared to pied flycatchers breeding further north (Sanz & Moreno, 2000). If migration distance, and not solely the latitudinal variation, was the main driver of telomere dynamics across populations, northern populations are expected to have shorter telomeres only among adults, as juveniles have not experienced any costs of migration yet. Furthermore, pied flycatcher populations across the breeding range are genetically differentiated from each other to some extent. Birds breeding in England, and in mountainous habitats in Spain and central Europe show the most differentiation to the extent that the Spanish birds are considered to be a separate subspecies of the pied flycatcher (*F. hypoleuca iberiae*) (Clements et al., 2021; Haavie, Sætre, & Moum, 2000; Lehtonen et al., 2012; Lehtonen et al., 2009). These genetic differences could create among-population differences in telomere length and dynamics. European pied flycatcher populations also differ in breeding dates, clutch size, number of fledglings (Sanz, 1997), and various egg characteristics (Morales et al., 2013; Ruuskanen et al., 2011), which might ultimately influence individual population-specific telomere length trajectories.

Here, we studied large-scale variation in the telomere biology of pied flycatchers by sampling nestlings and adult birds from six breeding populations across a north-south gradient across Europe. We specifically examined 1) overall patterns of telomere length variation across populations and life stages, 2) associations between telomere length and migration distance, 3) early-life telomere dynamics and body mass growth, and 4) relationships between telomere length and body mass at different ages across populations. We predict that 1) adult birds would show more among population variation in telomere length than juveniles, and the variation could be related to latitudinal differences in migration distance, *i.e.*, increasing migration distance would be associated with shorter telomeres. We also predict that 2) nestling telomere shortening would be negatively related to nestling growth rate within populations due to high metabolic costs of growing and selective energy allocation to somatic growth, and 3) that this relationship between early-life telomere shortening and growth rate might differ between populations due to possible differences in environment, genetics, or both.

## Methods

### Study populations

Data for this study were collected during the 2019 breeding season from six different pied flycatcher populations along the south-north axis of the breeding range: Valsaín, central Spain (40°54’N, 4°01’W), La Hiruela, central Spain (41°04’N, 3°27’W), East Dartmoor, southern England (50°36’N, 3°43’W), Kilingi-Nõmme, southern Estonia (58°7’N, 25°5’E), Turku, southern Finland (60°25’N, 22°10’E), and Oulu, northern Finland (65°0’N, 25°48ʹE). All birds were breeding in nest boxes in study areas established several years before this study (Fig. S1).

### Sample collection

Between early May and early June pied flycatcher nests were monitored in each study area for laying date (pied flycatchers lay one egg per day), clutch size (typically 4-8 eggs) and hatching date (on average 14 days from the start of incubation). The nestling period from hatching to fledging is *ca.* 15-17 days. One random chick per nest was sampled at days 5 and 12 (hatching day = day 0). By day 12, most of the chicks’ structural growth is already complete and the mass gain has peaked and flattened (Lundberg & Alatalo, 1992), while sampling later than this may cause premature fledging. Additionally, the social parents (*i.e.* adult birds feeding the chicks) in each nest were caught and sampled when their chicks were around 10 days old. Approximately 60 birds per population (20 chicks, 20 females, and 20 males; see exact sample sizes in figure legends) were sampled. In each population, nests for sampling were selected along the hatching date gradient to standardize the effects of hatching date on studied parameters. All birds were weighed after blood sampling. In case the exact age of an adult could not be determined based on ringing information, the adults were aged either as a one-year-old or older based on feather characteristics (Svensson, 1992). Ultimately, all adults were categorized either as a one-year-old (young) or older (old).

The same blood sampling protocol including blood storage buffers was applied to all populations to eliminate differences in sample collection and storage, as this might affect the subsequent telomere measurements (Reichert et al., 2017). Blood samples (10-30 µl from adults and 12-day chicks, 10 µl from 5-day chicks) were collected by puncturing the brachial vein with a sterile needle and collecting the blood with a non-heparinized capillary tube. Blood was diluted with *ca.* 65 µl of PBS for storage. The samples were kept cold while in the field and stored at -20° C at the end of the day. All the blood samples were shipped to University of Turku on dry ice for DNA extraction and telomere length quantification.

### Laboratory analyses

All the laboratory work was conducted at the University of Turku by TK. Four months after sample collection, DNA was extracted from whole blood using a salt extraction alcohol precipitation method (Aljanabi & Martinez, 1997). Extracted DNA was diluted with BE buffer (Macherey-Nagel, Düren, Germany). DNA concentration and purity were quantified using ND-1000-Spectrophotometer (NanoDrop Technologies, Wilmington, USA; see Table S1 for population-specific results). DNA integrity was checked using gel electrophoresis (50 ng DNA, 0.8% agarose gel at 100 mV for 60 min using MidoriGreen staining) on 25 randomly selected samples and was deemed satisfactory (8 adult, 8 fledgling, and 9 5-d old nestling samples, 1-3 samples per age class per population). Samples were diluted to concentration of 2.5 ng/µl, aliquoted and stored in -20° C until telomere length assessment.

**Table 1.**
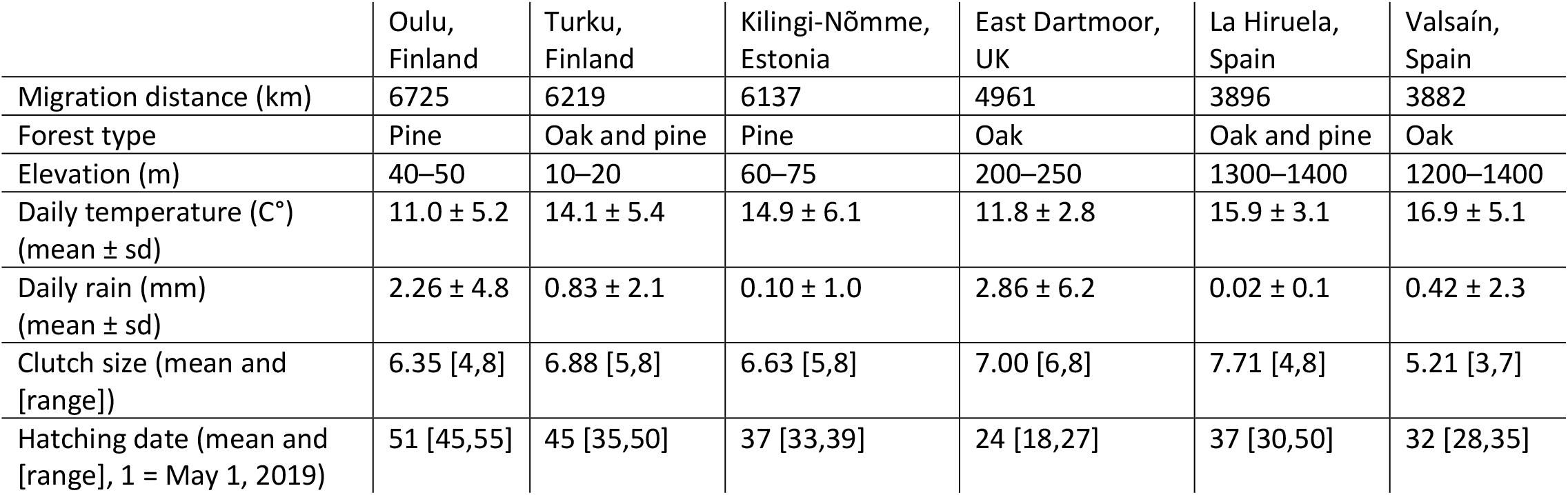
Differences in migration distance, forest type, elevation, mean daily temperature and rain during May-June 2019, clutch size, and laying dates among European populations of pied flycatcher. Migration distances were calculated at https://gps-coordinates.org/distance-between-coordinates.php, and elevations were estimated at https://en-gb.topographic-map.com/maps/s5d7/Europe/. Weather data were obtained from the following weather stations: Oulu: Oulunsalo Pellonpää weather station, Oulu, Finland; Turku: Artukainen weather station, Turku, Finland; Kilingi-Nõmme: Pärnu, Häädemees and Laadi weather stations, Pärnumaa, Estonia; East Dartmoor: Haytor weather station, Dartmoor, Devon, UK; La Hiruela: Colmenar viejo and Buitrago del Lozoya weather stations, Madrid, Spain; Valsaín: Segovia weather station, Castilia and Leon, Spain.

|  | Oulu,<br>Finland | Turku,<br>Finland | Kilingi-Nõmme,<br>Estonia | East Dartmoor,<br>UK | La Hiruela,<br>Spain | Valsaín,<br>Spain |
| --- | --- | --- | --- | --- | --- | --- |
| Migration distance (km) | 6725 | 6219 | 6137 | 4961 | 3896 | 3882 |
| Forest type | Pine | Oak and pine | Pine | Oak | Oak and pine | Oak |
| Elevation (m) | 40–50 | 10–20 | 60–75 | 200–250 | 1300–1400 | 1200–1400 |
| Daily temperature (C°)<br>(mean ± sd) | 11.0 ± 5.2 | 14.1 ± 5.4 | 14.9 ± 6.1 | 11.8 ± 2.8 | 15.9 ± 3.1 | 16.9 ± 5.1 |
| Daily rain (mm)<br>(mean ± sd) | 2.26 ± 4.8 | 0.83 ± 2.1 | 0.10 ± 1.0 | 2.86 ± 6.2 | 0.02 ± 0.1 | 0.42 ± 2.3 |
| Clutch size (mean and<br>[range]) | 6.35 [4,8] | 6.88 [5,8] | 6.63 [5,8] | 7.00 [6,8] | 7.71 [4,8] | 5.21 [3,7] |
| Hatching date (mean and<br>[range], 1 = May 1, 2019) | 51 [45,55] | 45 [35,50] | 37 [33,39] | 24 [18,27] | 37 [30,50] | 32 [28,35] |

Real-time quantitative PCR (qPCR) was used to assess relative telomere length, as previously described in birds (Criscuolo et al., 2009) and validated in the pied flycatcher (Kärkkäinen et al., 2019). qPCR quantifies the amount of telomeric sequence (T) relative to the amount of single copy gene sequence (SCG) resulting in relative telomere length (T/S ratio). Here, we used RAG1 as a SCG (verified as single copy using a BLAST analysis on the collared flycatcher *Ficedula albicollis* genome), as previously used in Kärkkäinen et al. (2020). Forward and reverse RAG1 primers were 5ʹ-GCAGATGAACTGGAGGCTATAA-3ʹ and 5ʹ- CAGCTGAGAAACGTGTTGATTC-3ʹ respectively, and forward and reverse telomere primers were 5ʹ- CGGTTTGTTTGGGTTTGGGTTTGGGTTTGGGTTTGGGTT-3ʹ (Tel-1b) and 5ʹ-GGCTTGCCTTACCCTTACCCTTACCCTTACCCTTACCCT-3ʹ (Tel-2b). Both primers were used at a final concentration of 200nM. For the qPCR assay, 5ng of DNA per reaction was used in a total volume of 10μl (8μl of master mix+2μl of DNA). The master mix contained 0.1μl of each primer, 2.8μl of water and 5μl of SensiFAST SYBR Lo-ROX master mix (Bioline, London, UK) per reaction.

Due to the closing down of many laboratory service providers in 2020 following the worldwide Covid-19 pandemic, the qPCR analyses were performed on two instruments. First, 71% of the samples were analyzed with QuantStudio™ 12 K Flex Real-Time PCR System (Thermo Fisher) using 384-well qPCR plates, while the rest of the samples were analysed with MicPCR (Magnetic Induction Cycler PCR Machine, Bio Molecular Systems) fitting 48-well plates. A subset of samples (n = 20) initially analysed with QuantStudio were rerun with MicPCR, and the technical repeatability between the two measurements was 0.851 (95% Cl [0.66, 0.94], P<0.001). The somewhat low agreement repeatability between the two machines stems mainly from the fact that the estimates obtained with MicPCR were consistently slightly higher than those estimated with QuantStudio (Fig. S2). The differences between the machines was controlled for by including qPCR plate ID, which consists of machine ID (QS or Mic) and a running number, as a random effect in the statistical models.

In QuantStudio the telomere and RAG1 reactions were run in triplicates adjacent to each other on the same plate. Each plate contained one golden sample that was run twice, one internal standard, and one negative control. The qPCR conditions were: an initial denaturation (1 cycle of 3min at 95°C), 40 cycles with first step of 10s at 95°C, second step of 15s at 58°C and third step of 10s at 72°C, with melting curve analysis in the end. In the MicPCR, the samples were run in duplicates and the telomere and RAG1 reactions were performed on separate plates. Each plate contained the golden sample twice (same as used with QuantStudio), and the internal standard. The qPCR conditions in the MicPCR were: an initial denaturation (1 cycle of 3min at 95°C), 25/40 cycles (telomere/RAG1) with first step of 5s at 95°C, and second step of 25s at 60°C, with melting curve analysis in the end. Repeated samples from the same chick and the samples from its parents were analysed on the same plate, and samples from different populations were evenly distributed between all the plates and machines. Altogether 8 plates were analysed with QuantStudio, and 8 plates + 4 plate reruns analysed with MicPCR.

LinRegPCR (Ruijter et al., 2009) was used to determine the baseline fluorescence, the qPCR efficiencies and the quantification cycle (Cq) values. To validate the use of our qPCR approach at the between-population level, we examined whether populations differed in control gene Cq, as well as qPCR efficiencies for both control gene and telomere assays (Table S2). Control gene Cq-values did not differ among populations (*F*_5, 528_ = 1.48, *p* = 0.20), but there was significant variation in both control gene and telomere assay efficiencies despite of differences being small (Control gene: *F*_5, 528_ = 8.11, *p* <.0001; Telomere: *F*_5, 528_ = 12.66, *p* <0001; Table S2). Thus, we added both efficiencies as covariates in all the analyses described below that used telomere length as dependent variable. As the inclusion of these covariates did not affect the any of the main results, they were removed from the final models to reduce model parameters. Thus, we are confident that our qPCR approach is valid for comparison of these six populations.

**Table 2.** Results of linear mixed models explaining the effects of Age class and Population on telomere length

| Independent variable | Telomere length |  |  |  |
| --- | --- | --- | --- | --- |
| | Estimate ± se | df<sub>num,dem</sub> | F / $\chi^2$* | P |
| Fixed effects |  |  |  |  |
| Intercept | 0.35 ± 0.16 | 66.97 | | |
| Age class |  | 2, 255.2 | 55.06 | < .0001 |
| Population |  | 5, 109.9 | 14.14 | < .0001 |
| Random effect |  |  |  |  |
| Nest box | 0.12 ± 0.04 | 1 | 12.65 | 0.0004 |
| ID | 0.08 ± 0.06 | 1 | 1.56 | 0.22 |
| qPCR plate | 0.24 ± 0.09 | 1 | 48.50 | < .0001 |
| Residual | 0.45 ± 0.06 | | | |

\*F-tests were used for significance tests of fixed effects, likelihood ratio tests ( $\chi^2$ ) with mixture distributions and one-sided p-values were used for random effects.

As such, relative telomere length (T/S ratio, hereafter telomere length) was calculated based on plate-specific efficiencies (mean ± s.d. efficiencies were 1.90 ± 0.02 for RAG1 and 1.89 ± 0.06 for telomere) using the mathematical model presented in Pfaffl et al. (2001). Technical repeatability based on triplicate measurements of telomere length was 0.957 (95% CI [0.951, 0.962], p<0.001), and inter-plate repeatability based on samples measured on more than one plate was 0.897 (95% CI [0.831, 0.932], p<0.001). Age-adjusted within-individual repeatability of telomere length in chicks was 0.373 (95% CI [0.217, 0.518], p<0.001), which is close to the average value found for qPCR studies (Kärkkäinen, Briga, Laaksonen, & Stier, 2021).

### Statistical methods

We used linear models, linear mixed models, and correlation analyses to study population differences in telomere length and chick growth. Telomere length values were standardized with z-transformation using scale()-function in R (v. 3.6.2, R core team 2019) prior to analyses for better general comparability of the results (Verhulst, 2020). Statistical analyses were conducted with SAS statistical software version 9.4 (SAS Institute, Cary, NC, USA). The models were estimated using restricted maximum likelihood (REML) and the Kenward–Roger method was used to calculate degrees of freedom of fixed factors, and to assess parameter estimates and their standard errors. Normality and heteroscedasticity assumptions were checked visually by plotting the models’ residuals (normal probability plot, histogram, boxplot, and linear predictor plot – results not shown).

We started by examining potential population differences in telomere length by fitting a model with telomere length as the dependent variable, age (5-day chick, 12-day chick, one-year-old adult, older adult), population and their interaction as explanatory factors, and nest ID, bird ID, and qPCR plate as random effects. However, as there was no significant difference in telomere length between the one-year-old and older adults (post hoc pairwise comparison *p* = 0.75), the adults were grouped together and the same model was run with life stage (5-day chick, 12-day chick, adult), population and their interaction as independent variables. As the interaction term was not significant (see Results), it was removed from the final model that included the main effects of life stage and population. These analyses were run also with datasets that included only the telomere length estimates obtained with QuantStudio or MicPCR to ascertain that the potential differences are not explained by the differences between the qPCR machines. To examine whether the potential population differences in telomere length were related to migration distance, we separately tested for the correlations between migration distance and adult telomere length, and migration distance and chick TL at day 12. Since all the individuals in a population have the same distance value, only the average TL values per population were included in these analyses. The Spanish populations breed so close to each other compared to other populations in sampled in this study (distance in straight line: La Hiruela – Valsaín: 53 km) that they were considered as one population in this analysis. Migration distance was calculated as a straight distance in km between breeding site and non-breeding site coordinates which were estimated for populations from Finland and England as a centre area in the data presented in Ouwehand et al. (2016) and Bell et al. (in review) (Table 1). Estonian birds were assumed to winter in the same areas as Finnish birds, and Spanish populations in the same areas as English birds (Fig. S1). As an alternative, migration distance was also estimated as an additive distance for each population (*i.e.*, adding the straight distance between populations A and B to the additive migration distance estimate of population A) that takes better into account the non-linear migration routes of especially the northern populations. The analyses gave similar results for both migration distance estimates, and for simplicity, straight distance estimates are further reported.

To examine patterns of chick growth and telomere dynamics between populations in more detail, we first fitted three models with chick body mass at day 5, body mass at day 12, and growth rate (body mass change between day 5 and day 12) as dependent variables and population as fixed effect in all the models. Additionally, the body mass change model included the initial body mass (day 5) as a covariate. These models were run also with clutch size as a fixed effect to test if variation in number of siblings affects mass growth of the chicks. Then, we fitted a model with chick telomere change value (change between day 5 and day 12) as the dependent variable, population as an explanatory variable, and qPCR plate as a random effect. Since the change in chick telomere length was calculated by subtracting day 5 measurement from day 12 measurement (thus negative values indicate telomere loss), regression to the mean was corrected by following the equations in Verhulst et al. (2013). As growth may influence telomere dynamics, we examined the effects of body mass on telomeres between populations. Each dependent telomere variable (day 5, day 12, and change) was tested with population and corresponding body mass as explanatory variables, first as main effects and then also including the interaction term, resulting in six models. The effect of clutch size on telomere length and dynamics was also tested but ultimately removed since it was never significant. qPCR plate was included as random effect in all the models. To identify whether the overall relationship between body mass and telomere length stems from within-population effects, between-population effects, or both, we created two new mass-variables for each existing mass-variable (day 5, day 12, and change, thus six new variables in total) following within-group centering approach as explained by van de Pol and Wright (2009) to separate the potential within-population effects from the between-populations effects. First, we calculated a group mean for each population to use as a variable capturing the between-populations effect. Then, the population mean was subtracted from each individual mass in the corresponding population, to create a variable that captures the within-population effects. These variables were used as fixed effects in three models with corresponding telomere-variable as the dependent effect and qPCR plate as a random effect.

## Results

While there were clear effects of both life stage and population on pied flycatcher relative telomere length (Table 2, Fig. 1, Fig S3), the general pattern of telomere dynamics with age did not differ significantly between populations (Life stage × Population: *F*_10, 317.5_ = 0.40, *p* = 0.95; Fig. 1). Post hoc pairwise comparisons adjusted with the Tukey-Kramer method revealed that telomeres gradually shortened from day 5 to adulthood (Fig. 1, all p <.0001). Additionally, pied flycatchers from England (East Dartmoor) had significantly longer telomeres than any other population across life stages (all p <.0004) and both Spanish populations (Valsaín and La Hiruela) had significantly longer telomeres than the Estonian population (Kilingi- Nõmme, all p <0.027), and both Finnish populations (Turku and Oulu), but only before the p-value adjustments (all p < 0.028, except Valsaín-Turku p = 0.058; Fig. 1;). Similar results were obtained from the datasets that included only the samples analysed with QuantStudio or MicPCR (Table S3, Fig. S4) thus rest of the analyses were carried out with the entire dataset. There was no clear correlation between migration distance and relative telomere length, neither in adults (r= -0.63, p = 0.26, N = 5 populations) nor in fledglings (r = -0.69, p = 0.20, N = 5 populations) (Fig. S5).

**Figure 1.**
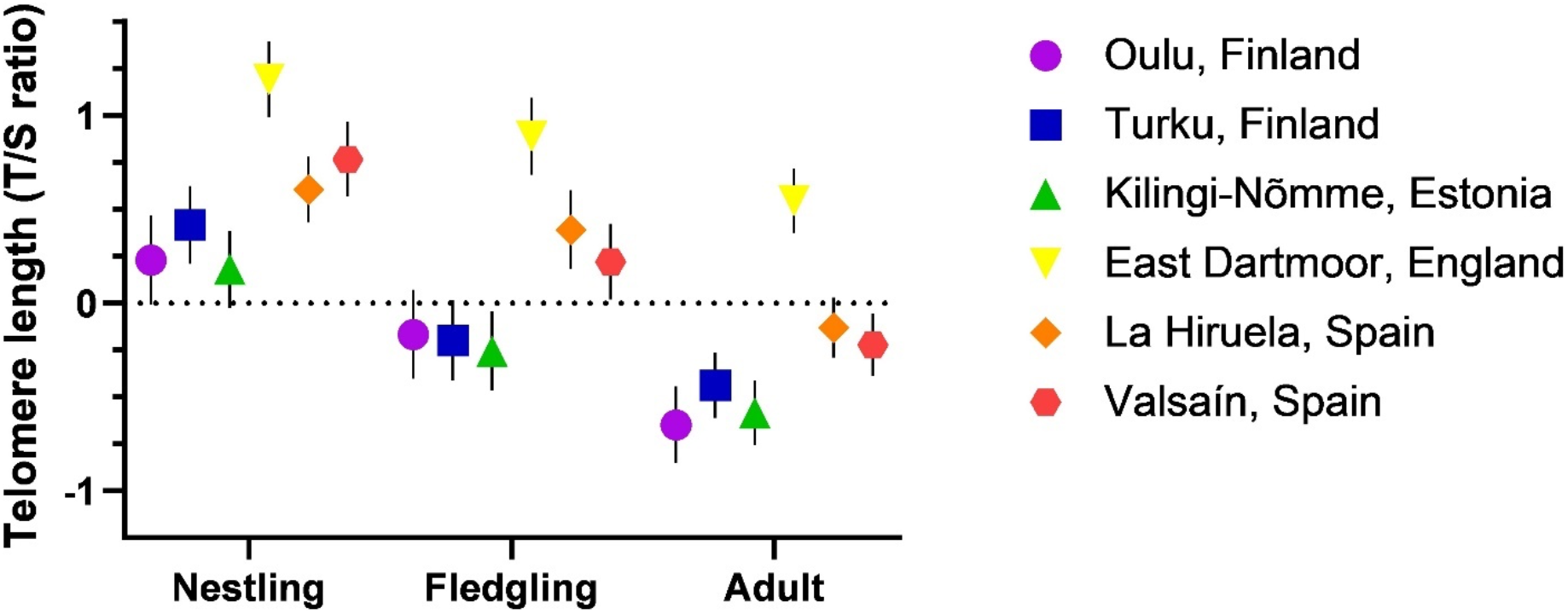
Relative telomere length in six pied flycatcher populations across a north-south gradient in Europe, from the early nestling period (Nestling; measured 5 days after hatching), to fledging (Fledgling; measured 12 days after hatching) and adulthood (Adult; measured at the end of their chicks’ rearing period). Values are estimated marginal means ± s.e.m based on z-scored telomere length values. Sample sizes [for Population: Nestling/Fledgling/Adult] are: Oulu: 19/19/41; Turku: 21/19/41; Kilingi-Nõmme: 22/20/43; East Dartmoor: 23/22/45; La Hiruela: 35/19/52; Valsaín: 24/23/49.

While focusing specifically on the early-life (nestling) period, the population of origin had a strong effect on body mass at day 5 (*F*_5, 133_ = 15.94, *p* <.0001; Fig. 2A), day 12 (*F*5, 116 = 2.53, *p* = 0.03; Fig. 2B) and growth rate (*F*_5, 105_ = 2.59, *p* = 0.03; Fig. 2C). Overall, chicks from both Spanish populations were smaller at day 5 but gained more body mass between day 5 and 12 to reach a fledging body mass similar to the other populations (Fig. 2). At day 12 the only significant difference in body mass was between the lightest (Turku) and the heaviest (Valsaín) chicks. However, while the clutch size had no significant effect on chick body mass at day 5 (Population: *F*_5, 132_ = 13.73, *p* <.0001; Clutch size: β = -0.009 ± 0.14, *F*_1, 132_ = 0.46, *p* = 0.50), it did affect negatively body mass at day 12 (Population: *F*_5, 115_ = 1.55, *p* = 0.18; Clutch size: β = -0.43 ± 0.18, *F*_1, 115_ = 8.40, *p* = 0.005) and consequently body mass growth (Population: *F*_5, 104_ = 1.18, *p* = 0.33; Clutch size: β = -0.35 ± 0.16, *F*_1, 104_ = 4.88, *p* = 0.03), diminishing the population differences in mass growth and fledgling mass (Fig. S6). There was also an effect of the population of origin on telomere shortening rate, albeit non-significant (*F*_5, 86.35_ = 2.19, *p* = 0.062; Fig. S7). Chicks from England (East Dartmoor), Spain (only Valsaín population) and southern Finland (Turku) tended to have higher shortening rates than the three other populations, although no post hoc tests were conducted due to the non-significance of the main effect (Fig. S7).

**Figure 2.**
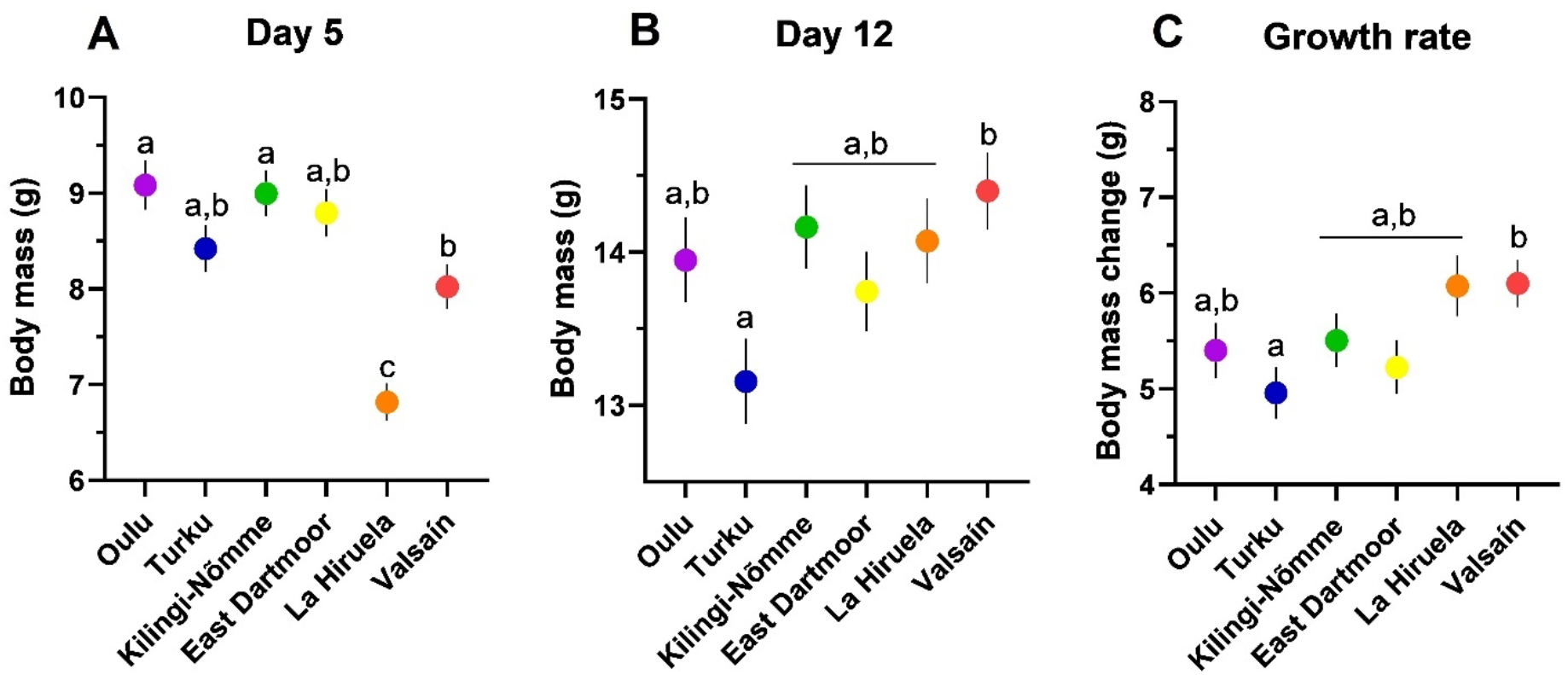
Pied flycatcher chick body mass at day 5 (A), day 12 (B) and growth rate (Δ mass between days 12 and 5; C) in six populations across a north-south gradient in Europe. Statistically significant differences after Tukey-Kramer adjustment for multiple comparisons are indicated with different letters. Values are estimated marginal means ± s.e.m. Sample sizes [for Population: Day5/Day12/Growth] are: Oulu, Finland: 19/19/17; Turku, Finland: 21/19/18; Kilingi-Nõmme, Estonia: 22/20/19; East Dartmoor, England: 20/22/18; La Hiruela, Spain: 33/19/18; Valsaín, Spain: 24/23/21.

Interestingly, while controlling for the population effect, chicks that were heavier at day 5 had shorter telomeres (β = -0.14 ± 0.06, *F*_1, 124.2_ = 4.92, *p* = 0.028) while there was no interaction between the population of origin and mass at day 5 in explaining telomere length at this age (*F*_5, 120.1_ = 0.77, *p* = 0.58). The population-centered model revealed that the observed association between telomere length and mass at chick day 5 was significant within-population (β = -0.14 ± 0.07, *F*_1, 129_ = 4.69, *p* = 0.03), but not significant between-populations (β = -0.11 ± 0.08, *F*_1, 123.8_ = 1.78, *p* = 0.18). Such a relationship was not significant by day 12, although the direction of the relationship remained similar (Mass at day 12: β = -0.09 ± 0.07, *F*_1, 114.6_ = 1.44, *p* = 0.23; within-population effect: β = -0.09 ± 0.08, *F*_1, 118.9_ = 1.16, *p* = 0.28; between-population effect: β = 0.005 ± 0.24, *F*_1, 129_ = 0.00, *p* = 0.99). Also, there was a significant interaction between population of origin and growth rate in explaining variation in telomere shortening rate (Population × Growth rate: *F*_5, 88.56_ = 2.36, *p* = 0.047). Specifically, fast growth was associated with faster telomere shortening in Finnish and Estonian populations, while the opposite or no relationship was found for English and Spanish populations (Fig. 3). Since the link between telomere change and body mass change varies among populations, there were no significant within- or between-populations effects in the relationship between telomere change and body mass change (within-population effect: β = -0.06 ± 0.06, *F*_1, 97.63_ = 1.17, *p* = 0.28; between-populations effect: β = -0.01 ± 0.09, *F*_1, 105_ = 0.03, *p* = 0.87).

**Figure 3.**
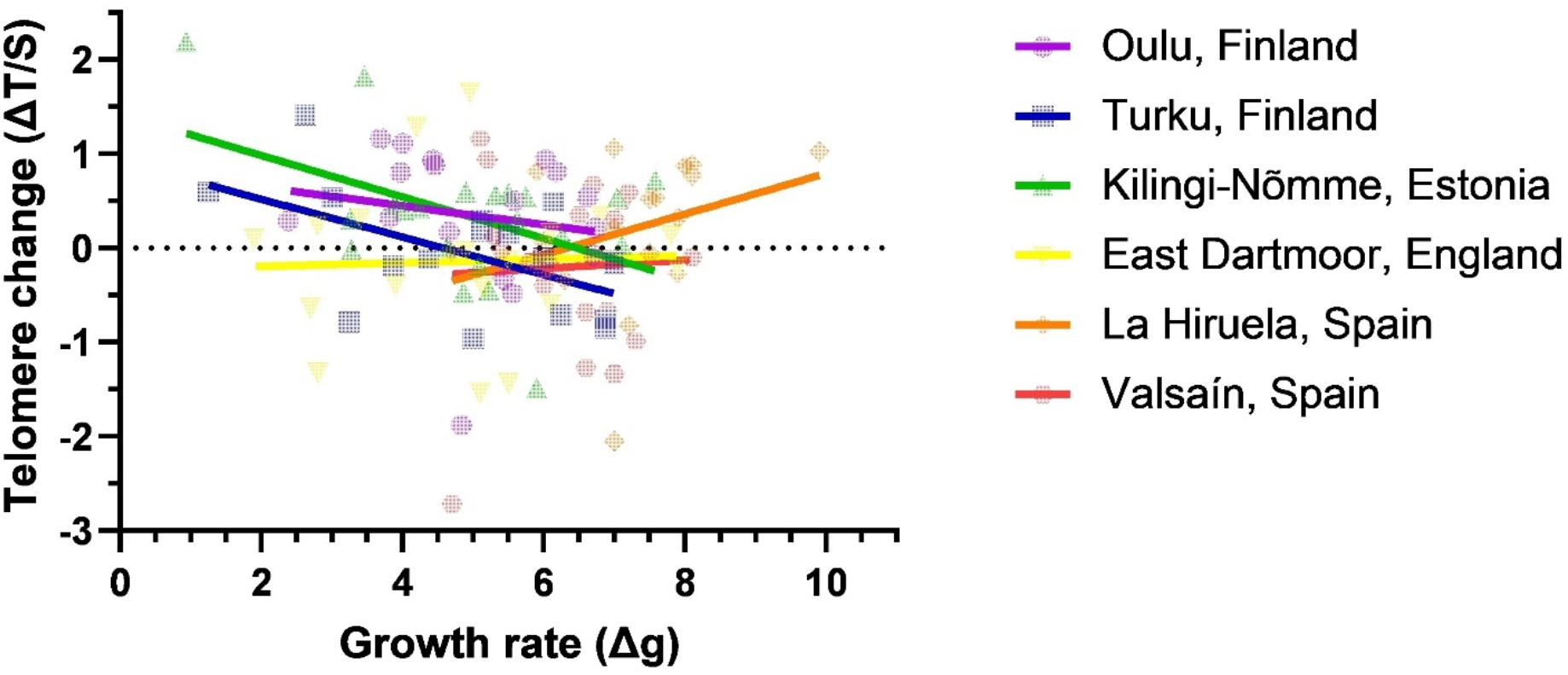
Association between growth rate (Δ mass between days 12 and 5) and telomere change (Δ telomere length between days 12 and 5 based on z-scored telomere length values) in pied flycatcher chicks in six populations across a north-south gradient in Europe. The interaction between population and growth rate was significant (*p* = 0.047) in explaining variation in telomere change (see results for details). Values are fitted with simple linear regression lines and individual data points are shown transparently for clarity. Sample sizes [for Population] are: Oulu: 17; Turku: 18; Kilingi-Nõmme: 19; East Dartmoor: 18; La Hiruela: 18; Valsaín: 21.

## Discussion

We found consistent variation in telomere length across European pied flycatcher populations and across different life stages (*i.e.* soon after hatching, close to fledging and in adulthood). There was no clear support for a relationship between migration distance and telomere length across populations. There was some indication that the rate of early-life telomere shortening varies between populations, but this effect was less pronounced than the pattern observed for chick body mass and growth rate. Heavier chicks had shorter telomeres in the early nestling period across all populations, an effect that was similar in direction but weaker close to fledging age. Interestingly, early-life growth rate was related to early-life telomere shortening rate, but in a population-dependent manner, with only northern populations exhibiting more telomere shortening when growing fast.

### Telomere dynamics across populations

As expected, telomeres shortened gradually both during early-life (-11.7%) and between fledging and adulthood (-12.6 %). Despite of some individual cases, we found no evidence for consistent telomere lengthening in any population. The overall dynamics observed with age did not differ between populations, despite variation in environmental conditions experienced across the North-South breeding range (Lundberg & Alatalo, 1992; Samplonius et al., 2018). Yet, there were clear differences in telomere length between populations. English birds (East Dartmoor) had the longest telomeres followed by Spanish birds (Valsaín and La Hiruela) having similar telomere lengths while the Estonian and Finnish birds (Kilingi- Nõmme, Turku, and Oulu) had the shortest telomere length. Notably, telomeres at the population level were not associated with increasing migration distance as birds breeding in the mid longitudinal part of the breeding range (England, East Dartmoor) had longer telomeres at any stage than the birds breeding further south. Furthermore, the pattern between migration distance and telomere length was similar in chicks at day 12. Our sample of different populations is clearly limited for deriving strong conclusions about this relationship. However, the English birds having the longest telomeres and the pattern of telomere change between nestling and adult stages being similar in all the populations indicates that the absolute differences in telomere length among populations are more attributable to other factors than to differences in migration distance. Previous studies have associated migratory lifestyle with shorter telomeres in dark-eyed juncos (*Junco hyemalis*) and longer migration distance with reduced fitness and survival in sanderlings (*Calidris alba*) (Bauer et al., 2016; Reneerkens et al., 2020). However, these associations might be more attributable to distinctive subspecies differences (migratory vs. resident populations; Bauer et al., 2016) and varying environmental conditions across distinct wintering sites (Reneerkens et al., 2020) rather than the migration distance *per se*, similarly as in Angelier at al. (2013). Nevertheless, due to logistical difficulties, our study is missing the pied flycatchers with the longest migration distance (breeding in west Siberia, around 1000 km longer migration route than Oulu population estimated with breeding site coordinates provided in Lehtonen et al. 2009) that could have been truly informative regarding this question. Especially considering that despite the long distances between populations, a population from western Siberia was not genetically differentiated from northern European populations (Finnish and Estonian), unlike the populations further south (English and Spanish) (Lehtonen et al., 2012; Lehtonen et al., 2009). Closer examination of other potential factors affecting population telomere length and inclusion of more populations are therefore needed to further ascertain our results (Burraco et al., 2021).

As there was no consistent link between telomere length and migration distance, correspondingly, telomere length did not show straightforward latitudinal variation coinciding with the pace of life -hypothesis either. While many life-history traits do show consistent latitudinal variation, in this study the latitudinal gradient can be disrupted by the mountainous habitat of the pied flycatchers breeding in the lowest latitudes as often, but not always, the effect of increasing elevation is similar to the effect of increasing latitude (Hille & Cooper, 2015). Alternatively, our latitudinal gradient was not extensive enough to show the possible effect on intraspecific telomere length (but see Kirby et al., 2017). Indeed, the biggest differences in trait variation with increasing latitude are observed in interspecific studies between tropical and temperate species or subspecies, which experience marked differences in *e.g.*, seasonal changes in food availability, that is often used to explain the occurrence of latitudinal variation (McNamara et al., 2008). As all the populations in this study are migratory, seasonality effects among populations are likely minimal.

There are latitudinal variation also in predator abundance and parasite prevalence of passerine birds in Europe (Díaz et al., 2013; Scheuerlein & Ricklefs, 2004), both of which can have a negative effect on telomeres (Asghar et al., 2015; Kärkkäinen et al., 2019), but these are unlikely explanations for our results. Predator abundance decreases with increasing latitude (Díaz et al., 2013), while we observed that birds from high northern latitudes (Estonia and Finland) had the shortest telomeres, which is the opposite of the expected predator effect. Similarly, prevalence of certain blood parasites was lowest in low latitudes increasing with increasing latitude (Scheuerlein & Ricklefs, 2004) but we observed the English birds to have longer telomeres than the Spanish birds. Instead, the observed latitudinal differences in telomere length might reflect the local environmental conditions *e.g.*, forest type, similarly as discussed by Quirici, Guerrero, Krause, Wingfield and Vásquez (2016). Deciduous forests of southern England might be more favourable breeding grounds than northern, conifer-dominated or southern montane forests, as also indicated by bigger clutches in mid-European latitudes compared to northern and southern populations (Sanz, 1997). Deciduous forests are also characterized by higher good-quality prey abundance than conifer-dominated forests (Burger et al., 2012), and good-quality prey might enable better telomere maintenance. Furthermore, egg yolk carotenoid levels are highest in central European pied flycatcher populations relative to southern and northern populations, although a population in Spain showed high concentrations of a few carotenoids (Eeva et al., 2011). Carotenoid concentrations in the eggs are reflective of female diet during egg laying (Török et al., 2007), and thus can be an indicator of environmental quality. Also, carotenoids work as antioxidants that alleviate oxidative stress (Surai, Fisinin, & Karadas, 2016) and possibly telomere shortening (Kim & Velando, 2015; Pineda-Pampliega et al., 2020). Therefore, possible higher levels of carotenoids in the diet of English (East Dartmoor), and to some extent the Spanish birds might contribute to the telomere length differences between populations we observed.

Genetic differences between pied flycatcher populations means that we cannot exclude that the average telomere length of a population would be genetically determined. In this study, population telomere lengths could be divided in three groups: Spanish (both Spanish populations), English (the English population), and the northern group (Estonian and both Finnish populations). The same distinction between populations can be done based on genetic differentiation as demonstrated by Lehtonen et al. (2012; 2009), who observed the Spanish and the English pied flycatchers to be genetically differentiated from each other and from the northern European populations, while the northern European populations could not be distinguished when using neutral genetic markers. Additionally, chromosomes contain non-terminal telomeric repeat sequences (interstitial telomeres, ITS) that are included in the relative telomere length measure (Foote, Vleck, & Vleck, 2013). Amounts of ITS might differ between populations, which could potentially explain why telomere length, but not shortening rate, differed markedly between populations.

### Early-life telomere dynamics and growth

The rate of nestling telomere shortening differed between populations but was not consistent along the north-south gradient, as the chicks from southern Finland (Turku), England (East Dartmoor), and one Spanish population (Valsaín) tended to have higher rates of telomere shortening than chicks from northern Finland (Oulu), Estonia (Kilingi-Nõmme), and the other Spanish population (La Hiruela). Curiously, those chicks growing in pine forests seemed to suffer less telomere shortening than those in oak forests, but this observation would require further testing using more replicates from different habitats. Since our data were collected over a single breeding season, we cannot exclude that the observed differences might simply reflect the local breeding conditions of the year. Typically, cold and rainy weather is not beneficial for the breeding of the pied flycatcher (Selonen et al., 2021). However, chicks from East Dartmoor, England, the rainiest and second coldest location in this study, grew as well as and showed longer telomeres than other chicks. Thus, more research is needed to evaluate the potential geographical variation in early-life telomere shortening and its underlying factors (Burraco et al., 2021).

Differences in chick growth between populations were clearer than differences in telomere shortening. We found that chicks from Spanish populations were lighter at day 5 but showed the highest growth rates from day 5 to day 12, and eventually matched the masses of chicks from other populations by day 12. This later growth peak in the Spanish flycatchers might be explained by elevation differences between populations. While all other populations in this study were at relatively low elevations (10-300 m above sea level), the Spanish flycatchers breed around 1 200 m above sea level. A previous study demonstrated great tit (*Parus major*) chicks, a species commonly breeding at low elevations, showed slower growth at high elevations (Stier et al., 2016), a difference potentially explained by the changes in prey availability, *i.e.* insect communities, with increasing elevation (Hodkinson, 2005). Chicks from bigger clutches gained less weight during days 5 and 12 and consequently were somewhat lighter at day 12, but this was not surprising considering that bigger clutch usually increase sibling competition that might negatively affect nestling growth (Nilsson & Gårdmark, 2001). On the contrary, telomere dynamics was not dependent on clutch size.

We found that, overall, at day 5, heavier chicks had shorter telomeres, and the tendency was the same at day 12. Closer examination revealed that this effect was significant within populations, *i.e*, heaviest chicks in each population also had the shortest telomeres in that population, but not between populations. However, close similarity of the within- and between-populations estimates (β = -0.14 vs. β = - 0.11) suggests that the effect might also be similar among populations, *i.e.*, populations whose chicks were the heaviest at day 5 also had the chicks with the shortest telomeres at same age. Indeed, previous studies have associated fast growth with faster telomere shortening (Monaghan & Ozanne, 2018; Stier, Metcalfe, et al., 2020; Tarry-Adkins, Martin-Gronert, Chen, Cripps, & Ozanne, 2008), as growth requires pronounced metabolic activity and cellular proliferation (Monaghan & Ozanne, 2018). Interestingly, chick growth affected more negatively telomere shortening in northern populations (Finland and Estonia) than in south-western ones (England and Spain). Similarly, telomeres of temperate juvenile stonechats (*Saxicola rubicola*) shortened during growth while those of tropical stonechats (*S. torquatus axillaris*) showed lengthening (Apfelbeck et al., 2019). However, to our knowledge, this sort of pattern has not been observed previously within one species. Chicks growing in mostly conifer-dominated forests further north might suffer from low-quality food (Burger et al., 2012), which together with the metabolic and oxidative stress caused by somatic growth could be detrimental for telomere maintenance. Additionally, carotenoids found in the eggs at mid latitudes, but also to some extent in southern Europe pied flycatcher populations (Eeva et al., 2011), might better safeguard chick telomeres as they grow (Min & Min, 2017; Pineda-Pampliega et al., 2020).

## Conclusion

To our knowledge, we provide the first study assessing large-scale geographical population differences in telomere length and dynamics (Burraco et al., 2021). Our results show that European pied flycatcher populations exhibit differences in mean telomere length both in chicks and adults, but that these differences do not vary consistently over latitudinal gradient. Instead, they might reflect more local environmental conditions and/or genetical differences. These marked population differences in telomere length dispute the common practice of using ‘species’ as unit in meta- and comparative analyses, as recently suggested by Canestrelli et al. (2020) and highlight the need to study telomeres at between-population level (Burraco et al., 2021). Future studies would benefit from closer examination of potential factors driving the observed between-population differences, and from assessing whether these differences in telomere length translate into between-population differences in lifespan, survival, and/or fitness proxies.

## Acknowledgements

We want to thank Angela Moreras for her invaluable help collecting the data, Carles Dura for ageing the adult birds in Oulu, and Natural England landowner of the study site in England, and finally the Subject Editor Prof. Simon Verhulst, and Dr Amos Belmaker and two other anonymous reviewers for their comments that considerably improved the manuscript. The study was financially supported by the Finnish Cultural Foundation Varsinais-Suomi Regional Fund, Turku University Foundation, Finnish Cultural Foundation (grants to TK), Turku Collegium for Science and Medicine (grant to AS), and Ministerio de Ciencia, Innovación y Universidades, MICINN, Spanish Government (postdoctoral grant IJC2018-035011-I and project no. PGC2018- 097426-B-C21 to AC and PID2019-104835GB-I00 to JMP). JMP was also funded by ARAID Foundation.

## Ethics

The collection of blood samples in Finland was licenced by the Animal Experiment Board in Finland (authorization license ESAVI/3021/04), in Estonia by the Animal Procedures Committee (licence no. 107) of the Estonian Ministry of Agriculture, in England by the Home Office (PPL 3003283), and in Spain by the Bioethics Committee of CSIC and Comunidad de Madrid (Valsaín: Ref. PROEX 128/19; La Hiruela: 10/106650.9/19) and Castilla-La Mancha (PR-21-2017).

## Author contribution

TK, AS and TL conceived the study following an original idea of JM. TK, MB, AC, JMP, JP, JM, RLT, and VT collected the blood samples. TK conducted the laboratory analyses with AS assistance. TK and AS analyzed the data. TK and AS wrote the manuscript with input from TL and other co-authors.

## Data accessibility

All the data used in this study is publicly available in Figshare (doi: 10.6084/m9.figshare.16940677) and can be accessed at https://figshare.com/s/0be81937376795cfd2b0.

## Supplemental Information

**Table S1.**
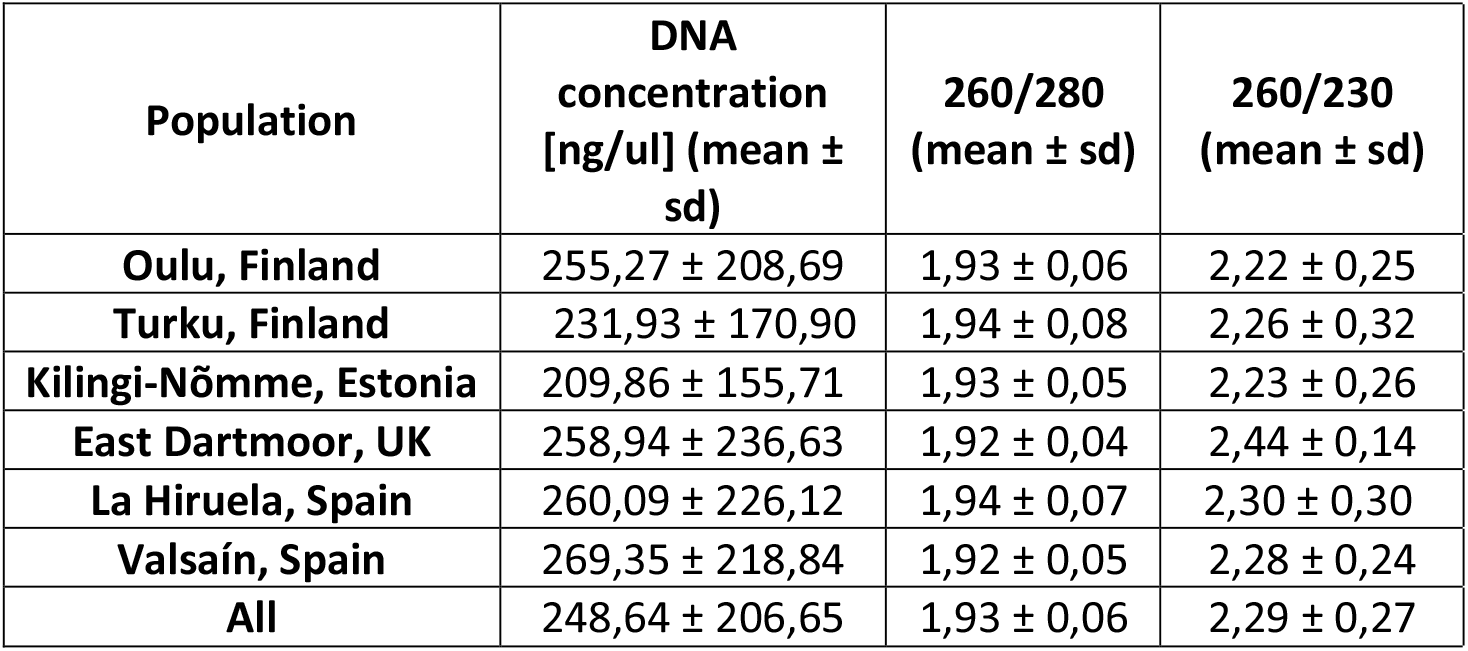
Results from DNA concentration and purity quantification using ND-1000- Spectrophotometer (mean ± sd). Large standard deviations for average concentration values are due to variation in tissue quantity among samples. All samples were however diluted to the concentration of 2.5 ng/µl before telomere length estimation. Three linear models (Concentration/[260/289]/[260/230] as dependent variable with Kenward-Roger approximation for degrees of freedom) were ran to test the differences among populations. Differences in DNA concentration were not statistically significant (F5, 531=1.02, p=0.40) while the differences in both 260/280 (F5, 531=2.45, p=0.03) and 260/230 (F5, 531=8.70, p<.0001) ratios reached statistical significance. Including the ratio-values as covariates in the statistical analyses presented in the main text with telomere length as dependent variable did not change the results or conclusion, thus these covariates were removed from the final models to reduce model parameters.

| Population | DNA concentration [ng/μl] (mean ± sd) | 260/280 (mean ± sd) | 260/230 (mean ± sd) |
| --- | --- | --- | --- |
| Oulu, Finland | 255,27 ± 208,69 | 1,93 ± 0,06 | 2,22 ± 0,25 |
| Turku, Finland | 231,93 ± 170,90 | 1,94 ± 0,08 | 2,26 ± 0,32 |
| Kilingi-Nõmme, Estonia | 209,86 ± 155,71 | 1,93 ± 0,05 | 2,23 ± 0,26 |
| East Dartmoor, UK | 258,94 ± 236,63 | 1,92 ± 0,04 | 2,44 ± 0,14 |
| La Hiruela, Spain | 260,09 ± 226,12 | 1,94 ± 0,07 | 2,30 ± 0,30 |
| Valsaín, Spain | 269,35 ± 218,84 | 1,92 ± 0,05 | 2,28 ± 0,24 |
| All | 248,64 ± 206,65 | 1,93 ± 0,06 | 2,29 ± 0,27 |

**Table S2.** Population specific (Mean ± sd) efficiencies and Cq-values for control gene (SCG) and telomere (TELO) assays. Three linear models (SCG Cq/SCG Efficiency/TELO Efficiency as dependent variable with Kenward-Roger approximation for degrees of freedom) were ran to test the differences among populations. Differences in SCG Cq-values were not statistically significant (F5, 528=1.48, p=0.20) while the differences in both SCG (F5, 528=8.11, p<.0001) and TELO (F5, 528=12.66, p<.0001) efficiencies reached statistical significance. Including both assay efficiencies as covariates in the statistical analyses presented in the main text with telomere length as dependent variable did not change the results or conclusion, thus these covariates were removed from the final models to reduce model parameters.

| Population | SCG Efficiency<br>[mean ± sd] | SCG Cq<br>[mean ± sd] | TELO Efficiency<br>[mean ± sd] | TELO Cq<br>[mean ± sd] |
| --- | --- | --- | --- | --- |
| Oulu, Finland | 1.87 ± 0.11 | 23.83 ± 1.50 | 1.88 ± 0.08 | 7.35 ± 1.05 |
| Turku, Finland | 1.91 ± 0.10 | 24.13 ± 1.44 | 1.93 ± 0.08 | 7.61 ± 0.92 |
| Kilingi-Nõmme, Estonia | 1.87 ± 0.11 | 23.82 ± 1.20 | 1.89 ± 0.07 | 7.40 ± 0.72 |
| East Dartmoor, England | 1.94 ± 0.04 | 23.98 ± 1.12 | 1.95 ± 0.06 | 7.03 ± 0.65 |
| La Hiruela, Spain | 1.90 ± 0.09 | 23.73 ± 1.64 | 1.89 ± 0.08 | 7.22 ± 1.12 |
| Valsaín, Spain | 1.88 ± 0.10 | 23.75 ± 1.06 | 1.91 ± 0.08 | 7.02 ± 0.75 |

**Table S3.**
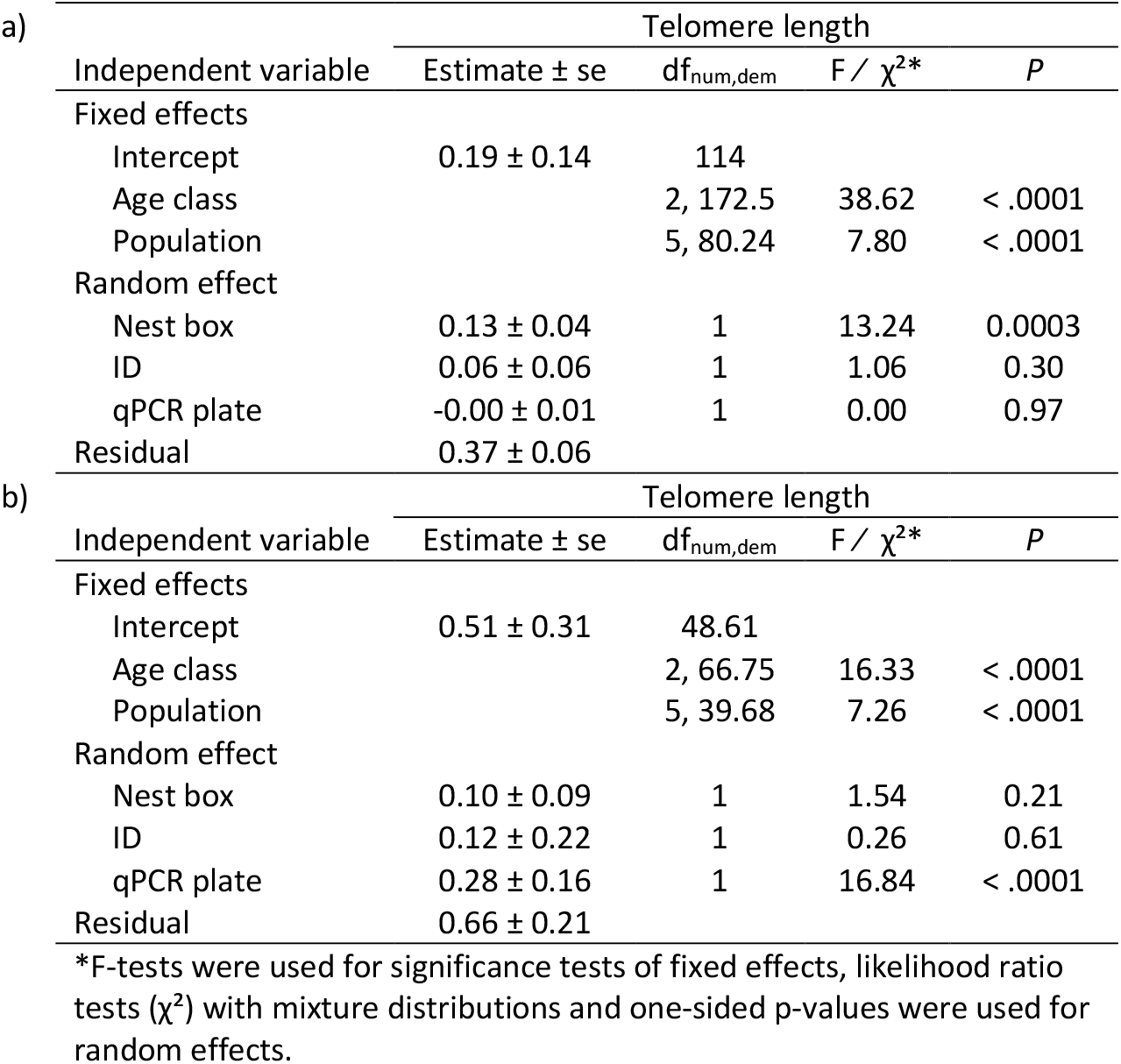
Results of linear mixed models explaining the effects of Age class and Population on telomere length using subsets of the whole data including samples analyzed only with a) QuantStudio or b) MicPCR

**Figure S1.**
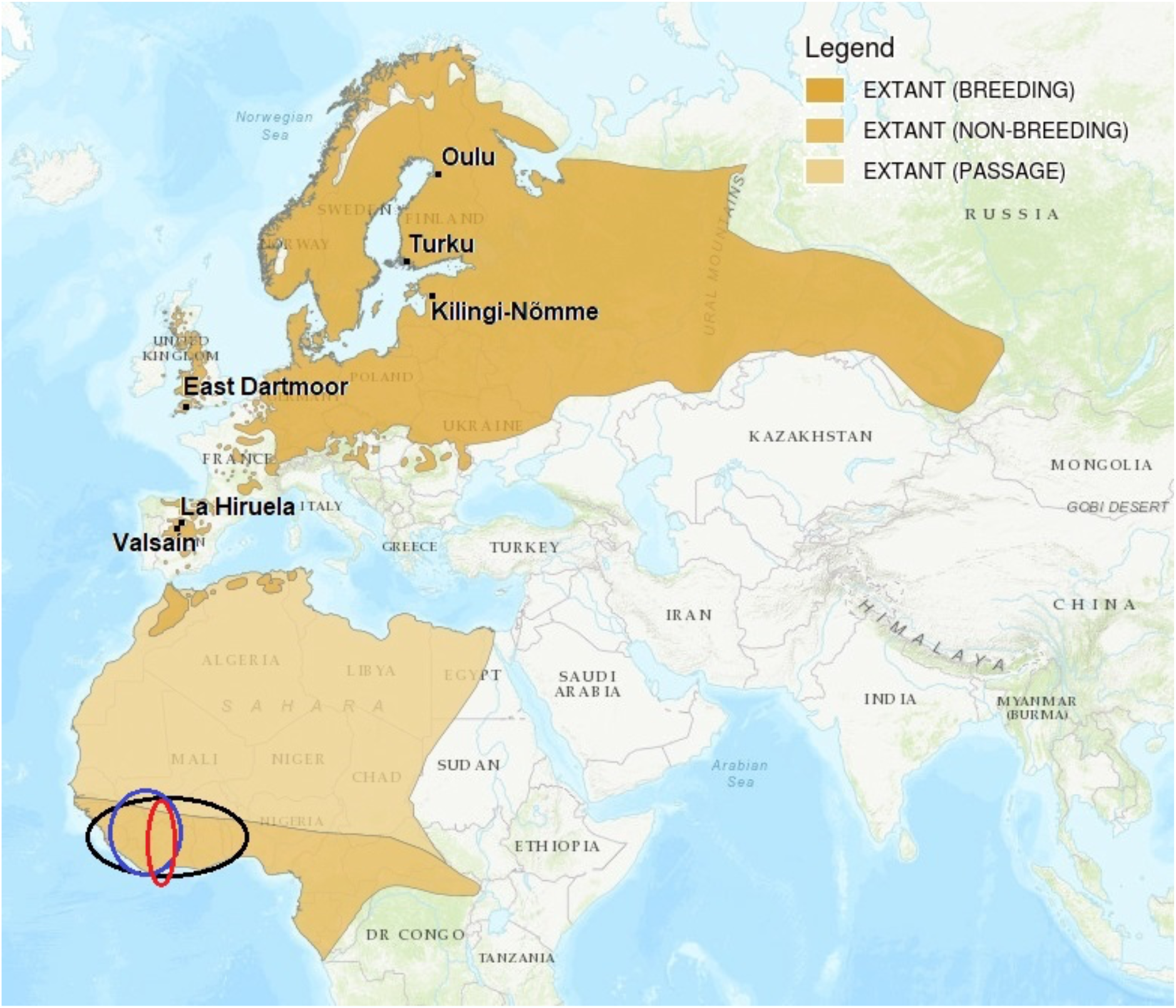
Locations of the study sites; breeding area of the pied flycatcher in Eurasia shown in orange. Birds from all populations are expected to migrate through Iberian Peninsula and west coast of Africa to their Sub-Saharan non-breeding grounds described in Ouwehand et al. 2016 (black circle; Finnish and Estonian birds blue circle; English and Spanish birds red circle). Map modified from: BirdLife International. 2018. Ficedula hypoleuca. The IUCN Red List of Threatened Species 2018: e.T22709308A131952521. https://dx.doi.org/10.2305/IUCN.UK.2018-2.RLTS.T22709308A131952521.en. Downloaded on 10 August 2021.

**Figure S2.**
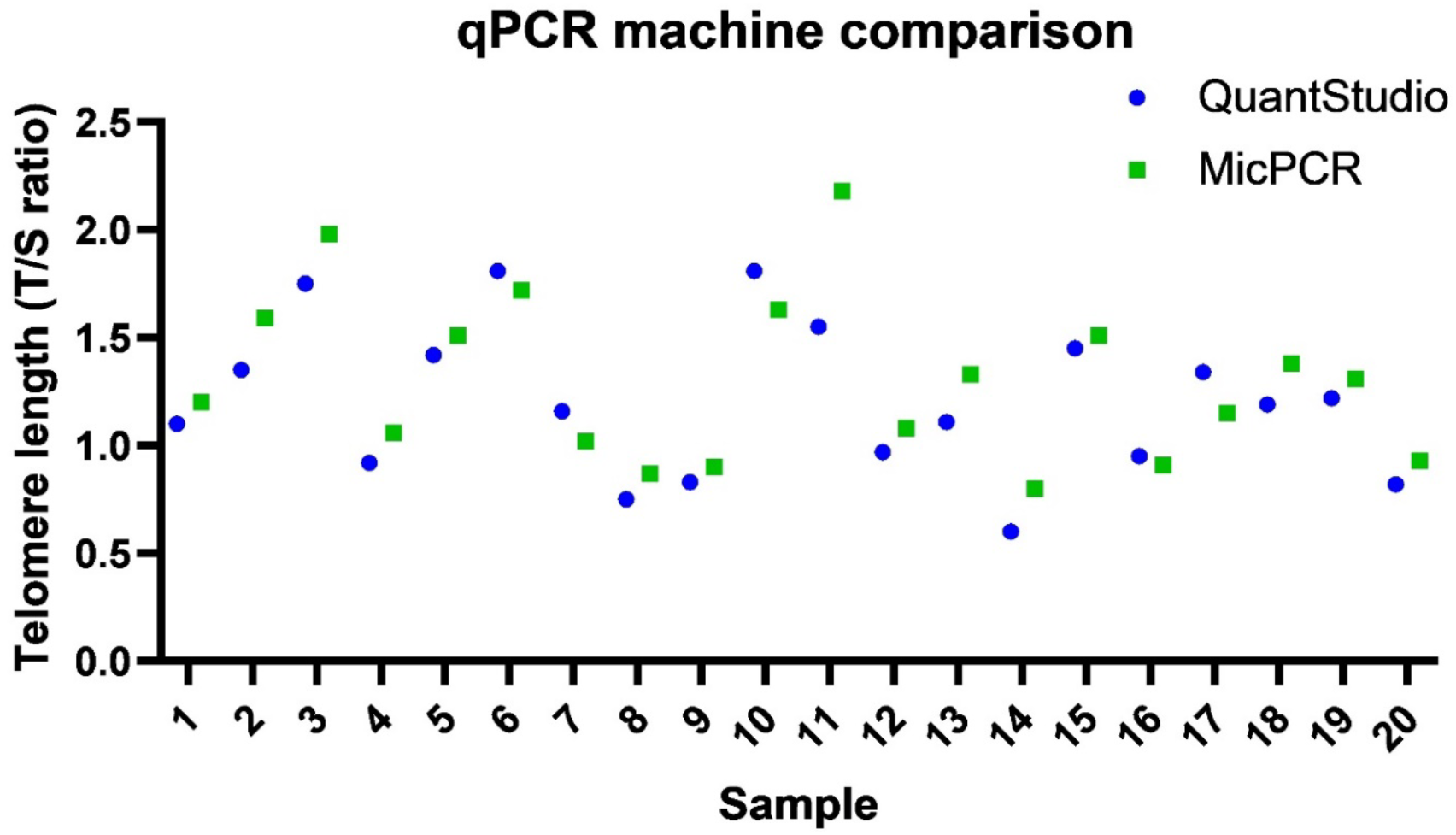
Illustrating the telomere lengths (T/S ratios) of the same sample measured both with QuantStudio and MicPCR. Telomere length estimates are consistently somewhat higher for MicPCR (15 out of 20 samples) accounting for somewhat low agreement repeatability of 0.851 (95% Cl [0.66, 0.94], P<0.001) between the two machines.

**Figure S3.**
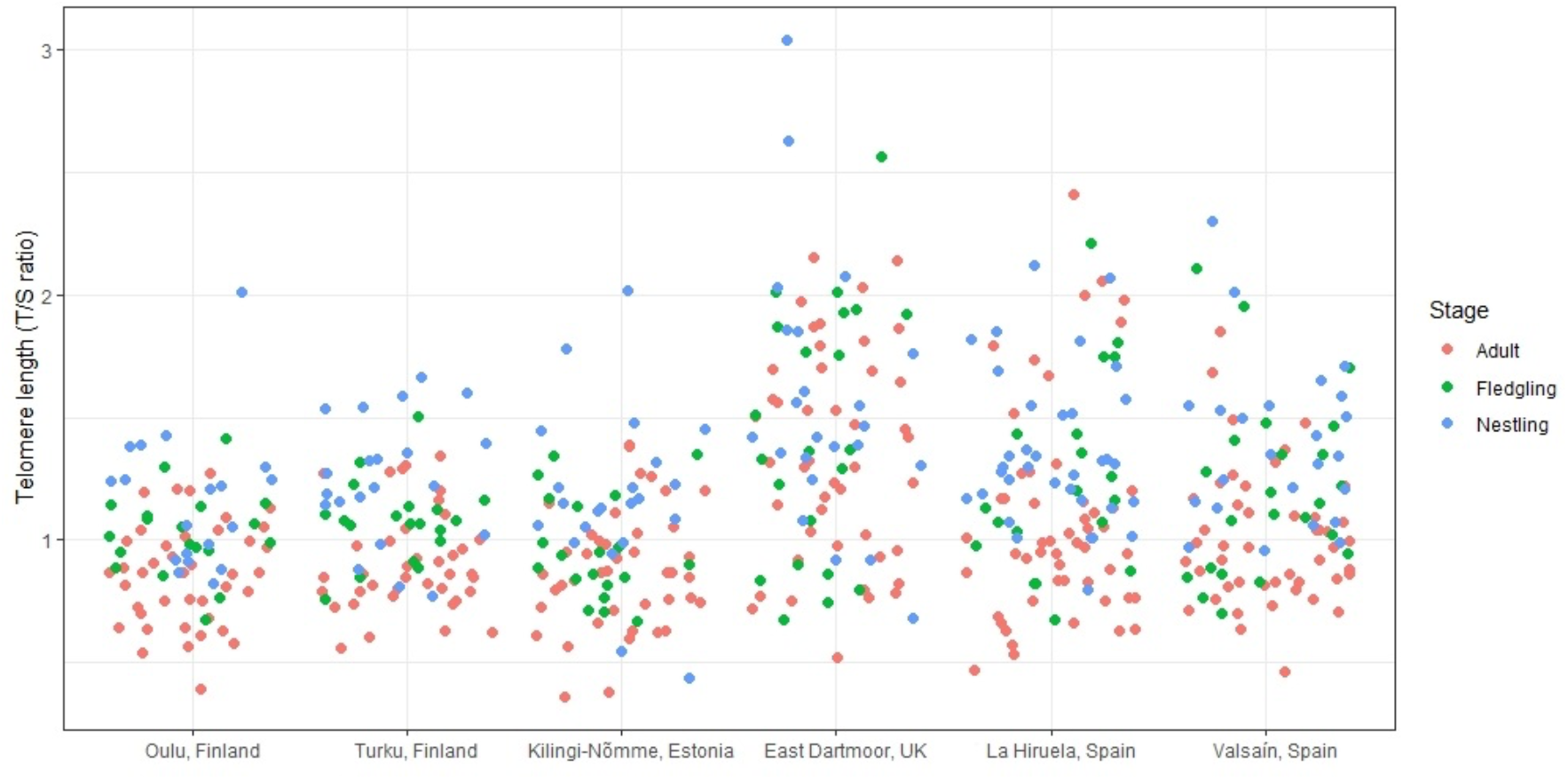
Individual raw telomere length values (T/S ratio) per population and age class. See sample sizes for Population: Nestling/Fledgling/Adult in the caption for Figure 1.

**Figure S4.**
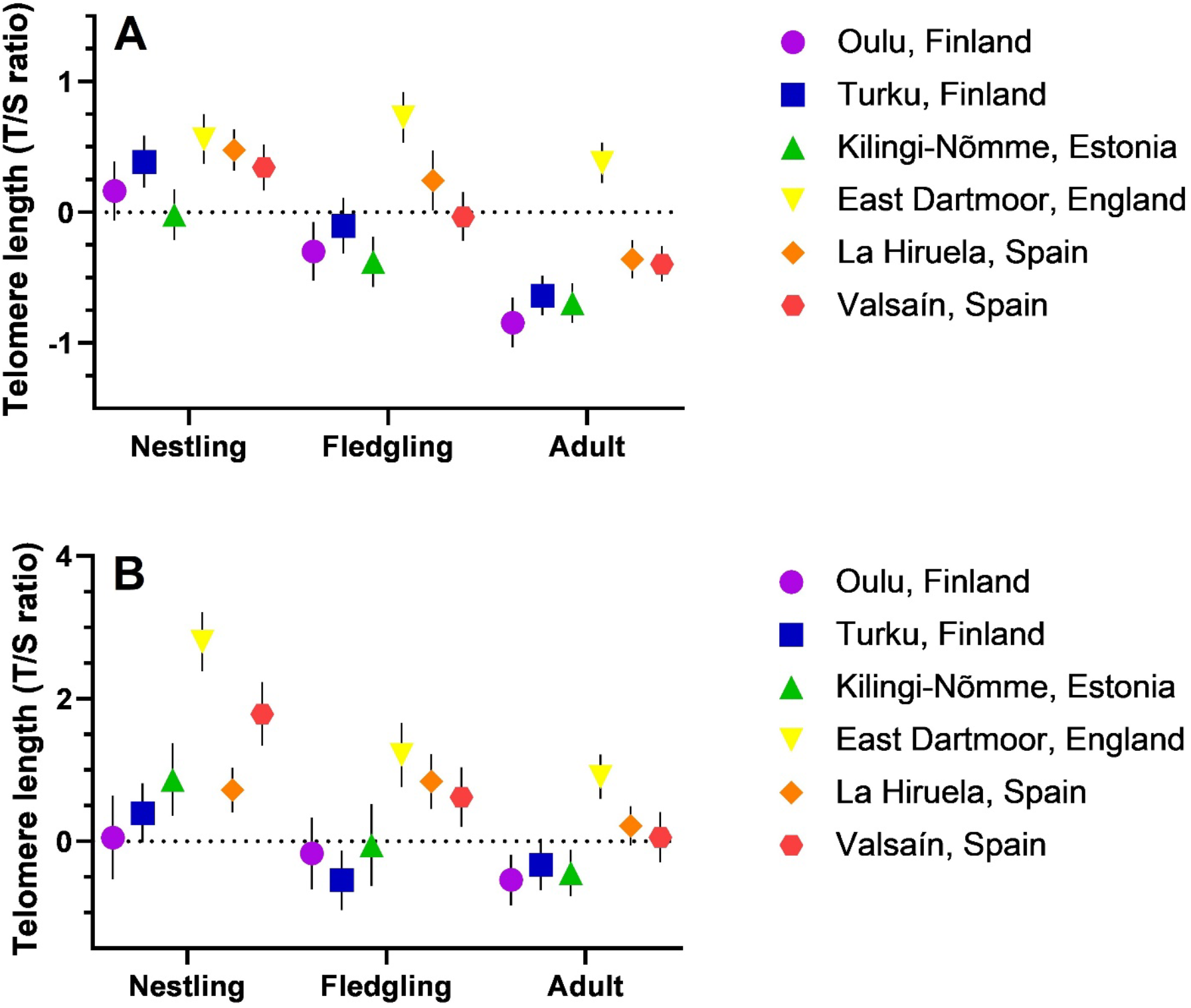
Relative telomere length in six pied flycatcher populations across a north-south gradient in Europe, from the early nestling period (Nestling; 5 days after hatching), to fledging (Fledgling; 12 days after hatching) and adulthood (Adult; end of the rearing period) using subsets of data including rTL values obtained only with a) QuantStudio, or b) MicPCR. Values are estimated marginal means based on z-scored telomere length values ± s.e.m. Sample sizes [for Population: Nestling/Fledgling/Adult] are a) Oulu: 16/15/29; Turku: 15/13/32; Kilingi-Nõmme: 18/17/31; East Dartmoor: 17/17/31; La Hiruela: 23/12/33; Valsaín: 19/17/39, and b) Oulu: 3/4/12; Turku: 6/6/9; Kilingi-Nõmme: 4/3/12; East Dartmoor: 6/5/14; La Hiruela: 12/7/19; Valsaín: 5/6/10.

**Figure S5.**
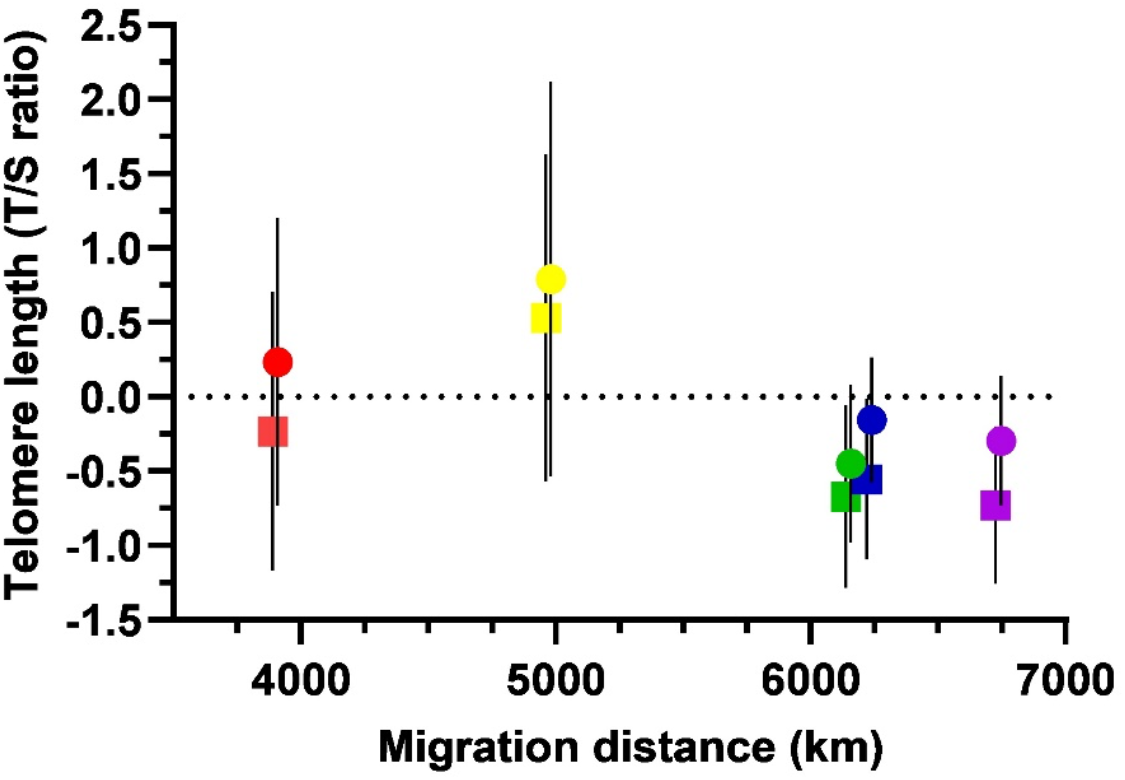
Associations between migration distance (km) and relative telomere length (mean based on z-scored values) in the pied flycatcher fledglings (12 days after hatching; circles) and adults (averaged breeding pair; squares). Standard errors of the means (± sem) have been added to illustrate the population variation in telomere length. Fledgling values (circles) have been moved slightly to the right to clarify the error bars. Populations from the shortest migration distance to the longest: Spain (average of Valsaín and La Hiruela, red), England (East Dartmoor, yellow), Estonia (Kilingi-Nõmme, green), southern Finland (Turku, blue), and northern Finland (Oulu, purple).

**Figure S6.**
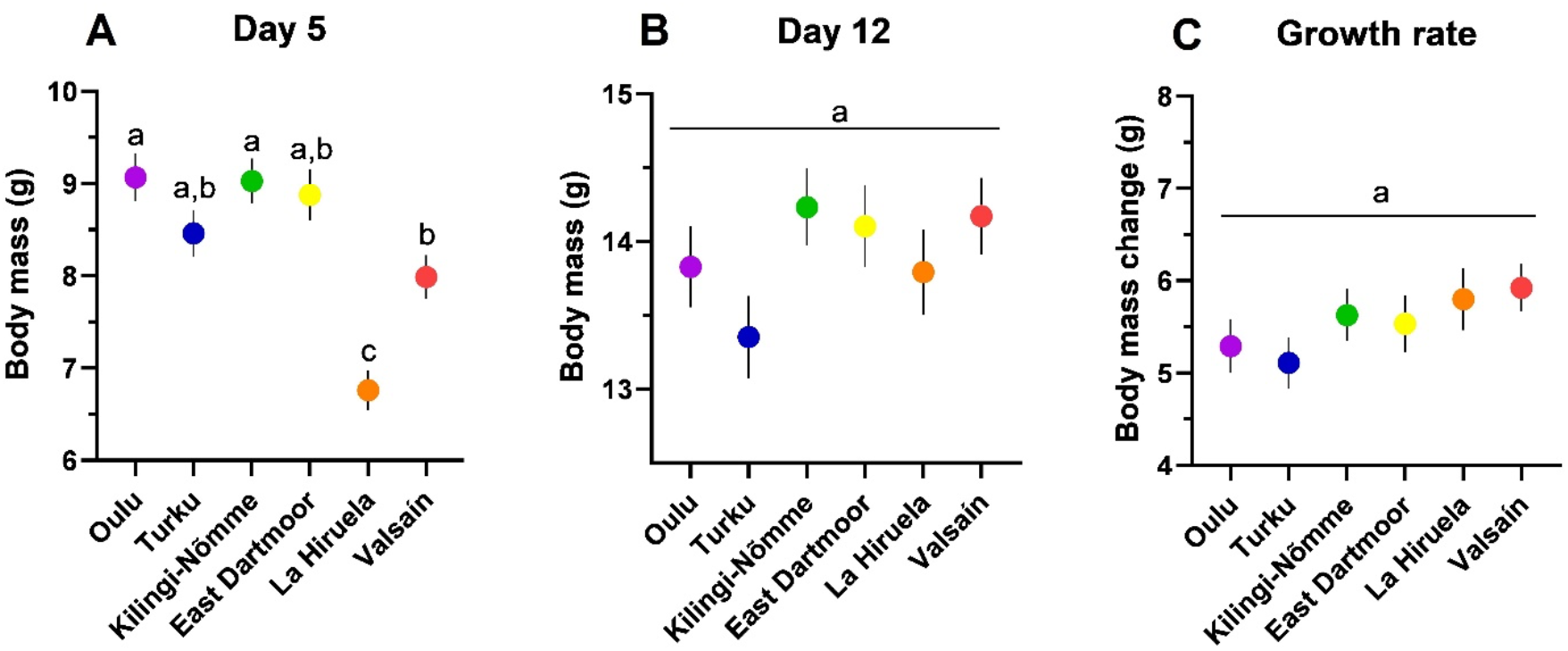
Pied flycatcher chick body mass adjusted for clutch size at day 5 (A), day 12 (B) and growth rate (Δ mass between days 12 and 5; C) in six populations across a north-south gradient in Europe. Statistically significant differences after Tukey-Kramer adjustment for multiple comparisons are indicated with different letters. Values are estimated marginal means ± s.e.m. Sample sizes [for Population: Day5/Day12/Growth] are: Oulu, Finland: 19/19/17; Turku, Finland: 21/19/18; Kilingi-Nõmme, Estonia: 22/20/19; East Dartmoor, England: 20/22/18; La Hiruela, Spain: 33/19/18; Valsaín, Spain: 24/23/21.

**Figure S7.**
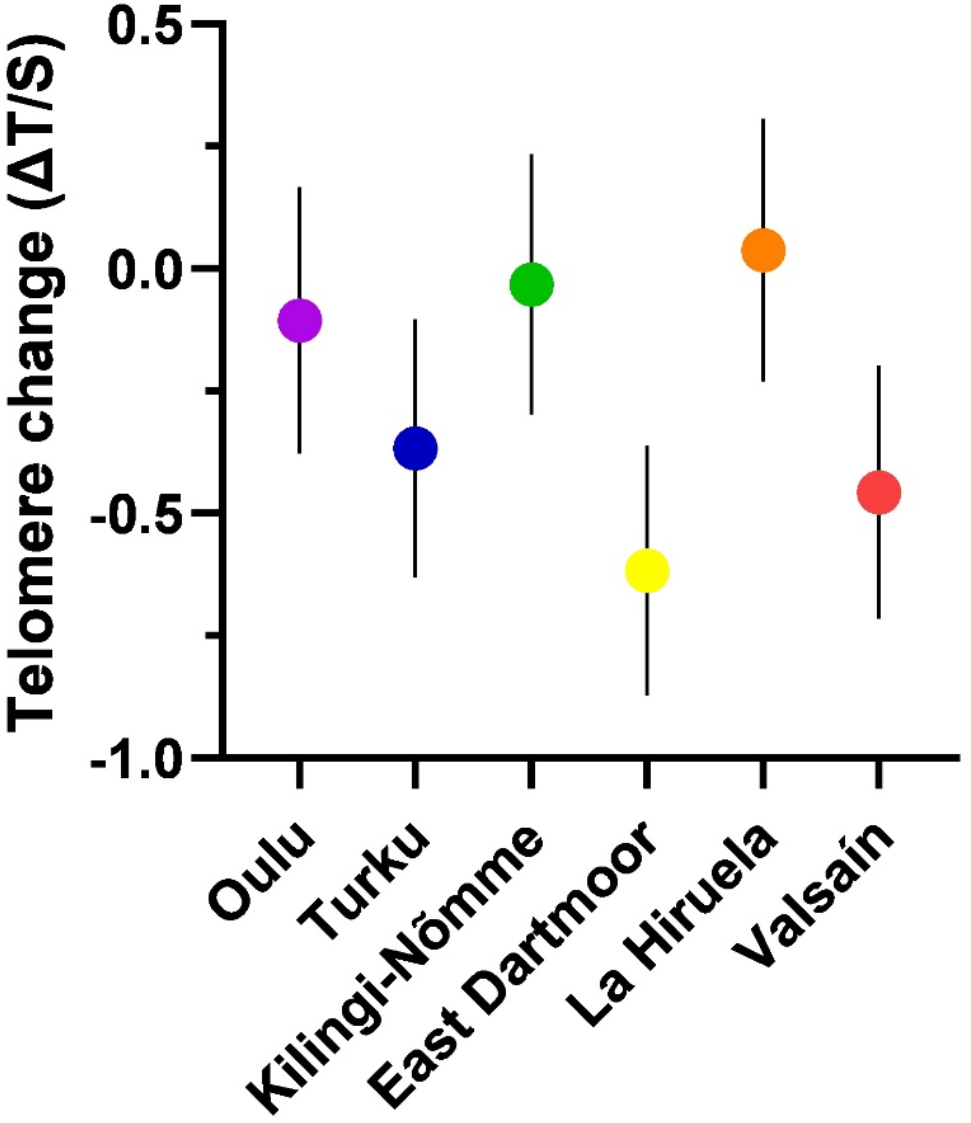
Change in relative telomere length during nestling period in the pied flycatcher (Δ telomere length between days 12 and 5) in six populations across a north-south gradient in Europe. The effect of population was marginally significant (*p* = 0.06) in explaining variation in early-life telomere change (see results for details). Values are estimated marginal means based on z-scored telomere length values ± s.e.m. Sample sizes [for Population] are: Oulu, Finland: 17; Turku, Finland: 18; Kilingi-Nõmme, Estonia: 19; East Dartmoor, England: 21; La Hiruela, Spain: 18; Valsaín, Spain: 21.

